# Identification of combinatorial colistin resistance mutations in *Shewanella algae*

**DOI:** 10.1101/2022.12.01.518641

**Authors:** Yao-Ting Huang, Yan-Chiao Mao, Chien-Hao Tseng, Chia-Wei Liu, Po-Yu Liu

## Abstract

**Background:** Colistin is one of the last-line antimicrobial agents against drug-resistant gram-negative bacteria. Currently, little is known about the genetic mechanisms underlying colistin resistance in *Shewanella algae*, partly due to complex epistatic interactions among multiple genes.

**Methodology/Principal Findings:** This study sequenced, assembled, and compared the genomes of 23 mcr-negative colistin-resistant *Shewanella algae* from marine, clam, oyster, and human. Comparative genomics and computational approach were applied to find combinatorial mutations. A combination of three mutations (PmrB451, PmrE168, PmrH292) was found to be strongly associated with colistin resistance in *Shewanella algae*.

**Conclusions/Significance:** This study demonstrates a computational approach for identifying epistatic-interacted mutations.

**Author summary:** *Shewanella algae* is an emerging pathogen related to Neglected Tropical Diseases (NTDs), including cobra-bite wound infections, marine injuries or ingestion of contaminated seafood. *Shewanella algae* is intrinsic resistant to various classes of β-lactams. Additionally, growing resistance to colistin in *mcr*-negative *Shewanella algae* further limits therapeutic options, especially in resource-limited regions. Currently, little is known about the genetic mechanisms underlying colistin resistance in *Shewanella algae*, partly due to complex epistatic interactions among multiple genes. We conduct comparative genomics to identify combinatorial colistin resistance mutations in *mcr*-negative colistin-resistant *Shewanella algae* and a combination of three mutations (PmrB451, PmrE168, PmrH292) is strongly associated with colistin-resistance.

## Introduction

Colistin is a critically important antimicrobial in humans with its efficacy against multi-drug resistant Gram-negative bacteria. However, rapid resistance towards colistin has been reported in diverse bacterial species globally. In addition, these bacteria are often co-resistant to other clinically important antimicrobials [1]. Resistance to colistin is partly caused by the phosphoethanolamine transferase enzymes, termed mobile colistin resistant (*mcr*) genes, transferred via mobile genetic elements [2, 3]. On the other hand, resistance of *mcr*-negative bacteria is mostly due to chromosomal mutations in two component systems (e.g., *PmrAB* or *PhoPQ*) [4]. These mutations often alter the outer membrane via addition of 4-amino-4-deoxy-l-arabinose (l-Ara4N) to phosphate groups of the LPS lipid A region, which reduces negative charge and decreases the acquisition of polycationic antimicrobial peptides and lipopeptide polymyxins [5, 6]. However, no consensus loci of chromosomal mutations leading to colistin resistance were found.

Antibiotic resistance-associated chromosomal mutations often come at the cost of fitness cost. For instance, mutations which reduce the influx of antibiotics by altering the outer membrane can decrease the absorption of nutrient compounds as well [7]. Similarly, overexpression of efflux pumps may also exporting beneficial compounds out of the cell [8]. Nevertheless, resistant strains may regain growth rate to wild-type levels by acquiring second or third mutations [9]. These deleterious/beneficial mutations can compensate for the fitness cost of each other [10]. Hence, chromosomal mutations often interact epistatically to reduce the overall fitness cost while developing antibiotic resistance [11, 12]. These complex epistatic interactions among multiple genes bring great challenges to the identification of colistin resistance genetic markers [5, 13].

*Shewanella algae* is an emerging pathogen associated with a wide range of diseases and serves as reservoirs of antibiotic resistance determinants in aquaculture. *S. algae* can be transmitted to human beings by water contact or ingestion of contaminated seafood. The organism was intrinsic resistant to various classes of *β*-lactams. In addition, resistance to colistin became common in *mcr*-negative *S. algae*, raise concern about the treatment failure and their environmental spread of colistin resistance [14]. To date, little is known about the genetic and molecular mechanisms underlying acquired resistance in *S. algae*, particularly for colistin.

In this study, we conducted comparative genomics and developed a computational approach to identify combinatorial colistin resistance mutations in *mcr*-negative colistin-resistant *Shewanella algae*. We compare single locus analysis with multiple combined mutations identified by novel combinatorial approaches. Mutations picked by single locus analysis showed significant association albeit with limited prediction accuracies. Whereas the integration of combinatorial approaches identified a combination of three mutations and achieves much higher prediction accuracies. Functional analysis of these three mutations revealed their potential impact on two-component systems related to outer membranes.

## Materials and Methods

### Isolates and antimicrobial susceptibility testing

*S. algae* isolates from different sources (Supplemental Table 1) were frozen with Luria Bertani (LB) broth containing 30% V/V glycerol and stored at −80°C until used as previous described [15]. Initial species identification was performed using Vitek 2 (bioMérieux, Inc., Durham, NC, USA) following the manufacturer’s instructions. The minimum inhibitory concentration values for colistin were determined by broth microdilution. Susceptibilities were interpreted using an MIC of 2 μg/mL as clinical breakpoint [16].

**Table 1.** Comparison of prediction accuracy of colistin resistance using different approaches. Comparison of prediction accuracy of colistin resistance using individual mutations and combined mutations identified by machine learning. The accuracies of all possible combinations of individual mutations found by single locus analysis (SLA) were also illustrated.

| Method | Mutation Locus | Accuracy | Sensitivity | Specificity |
| --- | --- | --- | --- | --- |
| SLA | PmrC214 | 73.91% | 0.714 | 0.778 |
| SLA | PmrC364 | 78.26% | 0.929 | 0.556 |
| SLA | PmrH330 | 69.57% | 0.786 | 0.556 |
| SLA | PmrC214, PmrC364 | 69.57% | 0.786 | 0.556 |
| SLA | PmrC214, PmrH330 | 56.52% | 0.643 | 0.444 |
| SLA | PmrC364, PmrH330 | 73.91% | 0.714 | 0.778 |
| SLA | PmrC214, PmrC364, PmrH330 | 65.22% | 0.643 | 0.667 |
| **Combinatorial** | **PmrB451, PmrE168, PmrH292** | **86.96%** | **0.929** | **0.778** |

### Identification of colistin resistance determinants

The genomes of 23 *S. algae* strains were collected as a local database (Supplemental Table 2). A multi-database approach was used to identify antibiotic resistance genes [17]. All the assembled sequences were BLAST [18] searched against the Comprehensive Antibiotic Resistance Database [19] and ResFinder [20] with a threshold of 1e−5. Queries with best hits were validated using BLAST searches against the Integrated Microbial Genomes & Microbiomes system with a 95% sequence identity threshold [21]. All search results were manually curated to ensure consistency of annotations among the different databases. If there were conflicting nomenclatures, the NCBI nomenclature would be used. Colistin-resistant genes reported in previous literatures were further extracted [22], including LpxA, LpxC, MexB, MexR, PhoP, PhoQ, PmrA, PmrB, PmrC, PmrE, PmrF, PmrH, and PmrK.

The protein sequences of colistin-resistant genes were concatenated separately for each strain. Subsequently, multiple sequence alignment of concatenated protein-sequences was performed by MEGA 7.0, revealing 647 mutated amino acid loci. Each of the mutated loci were tested for association with colistin resistance via Fisher’s exact test. The obtained *P* values were log transformed (-log_10_ *P*) and visualized as circular Manhattan plot by R.

### Combined mutations identified by solving minimum test collection problem

The candidate sequences of 647 mutated amino acid loci were reduced to a minimum test collection problem, which aims to find a minimum subset of loci capable of distinguishing between colistin-sensitive and colistin-resistant strains. As the minimum test collection is an NP-hard problem, we computed the approximate solutions by improving a traditional greedy algorithm. A traditional algorithm selects a mutated locus able to distinguish most pairs between sensitive and resistant strains. Then, another locus that can distinguish the remaining (undistinguished) pairs of sensitive/resistant strains is selected. The entire selection process is repeated until all pairs of sensitive and resistant strains can be distinguished, producing combined mutation subsets. To expand the search space, the entire process is further repeated ten times by always removing the best choice, second best choice, and so on, creating multiple approximate solutions. The best among them is selected as the final solution.

The individual mutated loci identified by Fisher’s exact test, all possible combinations among them, and the final solution of the minimum test collection, were features in the machine learning experiments. We compared the prediction accuracy using different features by classifying them with support vector machine (SVM). Leave-one-out cross validation is used to avoid overfitting.

### Functional analysis of combined mutations

The protein sequences of three genes containing combined mutations (PmrB, PmrE, and PmrH) were extracted and aligned against protein domain database InterPro [23]. Four and three domains were found in PmrB and PmrE, respectively, while no known domains were found in PmrH.

## Results

### Resistome analysis

Resistome annotation revealed presence of 211 to 248 antibiotic-resistant genes (ARGs) in each of the assembled genomes. Antimicrobial susceptibility tests indicated 14 strains are resistant to colistin (Supplementary Table S3).

### Individual colistin-resistant mutations analysis

Genes associated with colistin resistance were extracted and their translated protein sequences were concatenated after resistome annotations (see Method). Multiple sequence alignment of concatenated protein sequences revealed 647 loci with amino acid substitution. We analyzed each of these loci to identify mutations likely leading to colistin resistance (Fig 1). Three mutations, PmrC330, PmrC364, and PmrH330, are significantly correlated with colistin resistance (*P*=0.018-0.036, Fisher’s exact test). The three mutations were then each further tested for their predictive accuracy for colistin resistance (Table 1). By using Support Vector Machine (SVM) and leave-one-out cross validation (LOOCV), the results indicated that the predictive accuracy for colistin resistance of each mutation ranged from 69.5% to 73.9% (Sensitivity=0.71-0.92, Specificity=0.55-0.77, AUC=0.67∼0.74) (Table 2, S1 Fig.). We further constructed sets of double and triple combinatorial mutations but still failed to improve the accuracy. In fact, a worse result was even obtained when combining PmrC214 and PmrH330 for prediction (Accuracy=56.52%).

**Fig 1.**
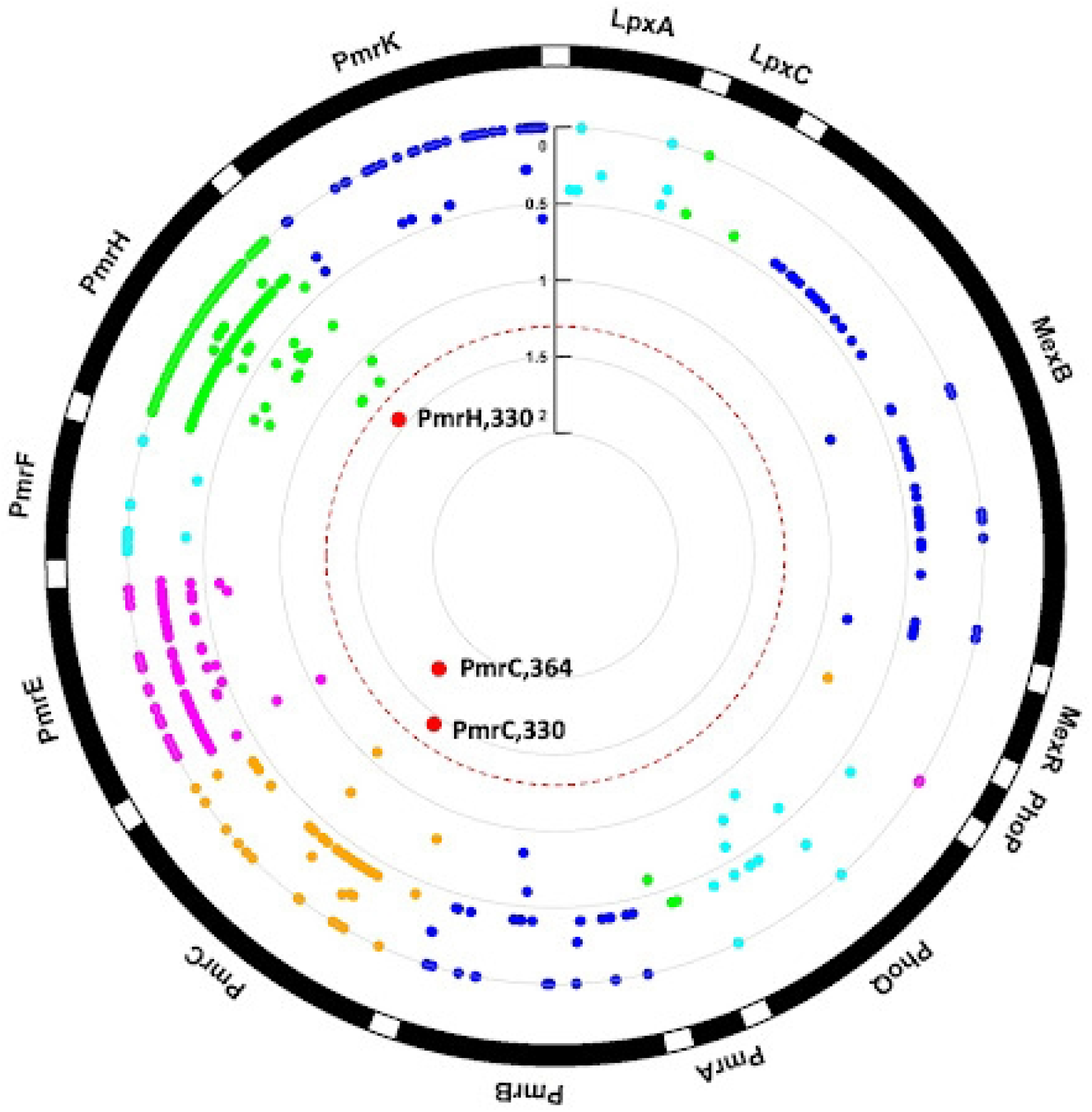
Circular Manhattan plot of amino acids association with colistin resistance. The *P* values of 647 mutations on 13 colistin-resistant genes are log transformed, whereas the red dashed circle represents *P*=0.05. Three mutations, PmrH330, PmrC364, and PmrC 330, are significantly associated with resistance.

### Combined resistant mutations found by combinatorial optimization

As colistin resistance is a complex synergistic effect caused by multiple genes in different pathways, single/individual mutation is insufficient to serve as genetic markers. To identify the combined effects of multiple mutations leading to colistin resistance, we employed the feature selection and classification of machine learning strategy (see Fig 2, Method). The feature selection stage solves a combinatorial optimization problem called minimum set cover, which aims to identify combinations of multiple loci capable of distinguishing resistance from sensitive strains. An SVM is trained by samples over these combined loci for prediction of strains with unknown phenotype. The selection algorithm uncovered a novel set of mutations (i.e., PmrB451, PmrE168, and PmrH292) completely different with previous loci (i.e., PmrC330, PmrC364, and PmrH330) in GWAS. We compared the prediction accuracy of combined mutations with three individual mutations found by single locus analysis (Table 2). The sensitivity, specificity, and accuracy of combined mutations were 92.9%, 77.8%, and 69.57%, respectively (Accuracy 86.96%, AUC 0.85), which are superior to both individual and combined mutations found in the single locus analysis (Accuracy 56.26 to 78.26%, AUC from 0.67 to 0.74). The results showed the combined effects of multiple markers were more strongly correlated with colistin resistance than individual mutations.

**Fig 2.**
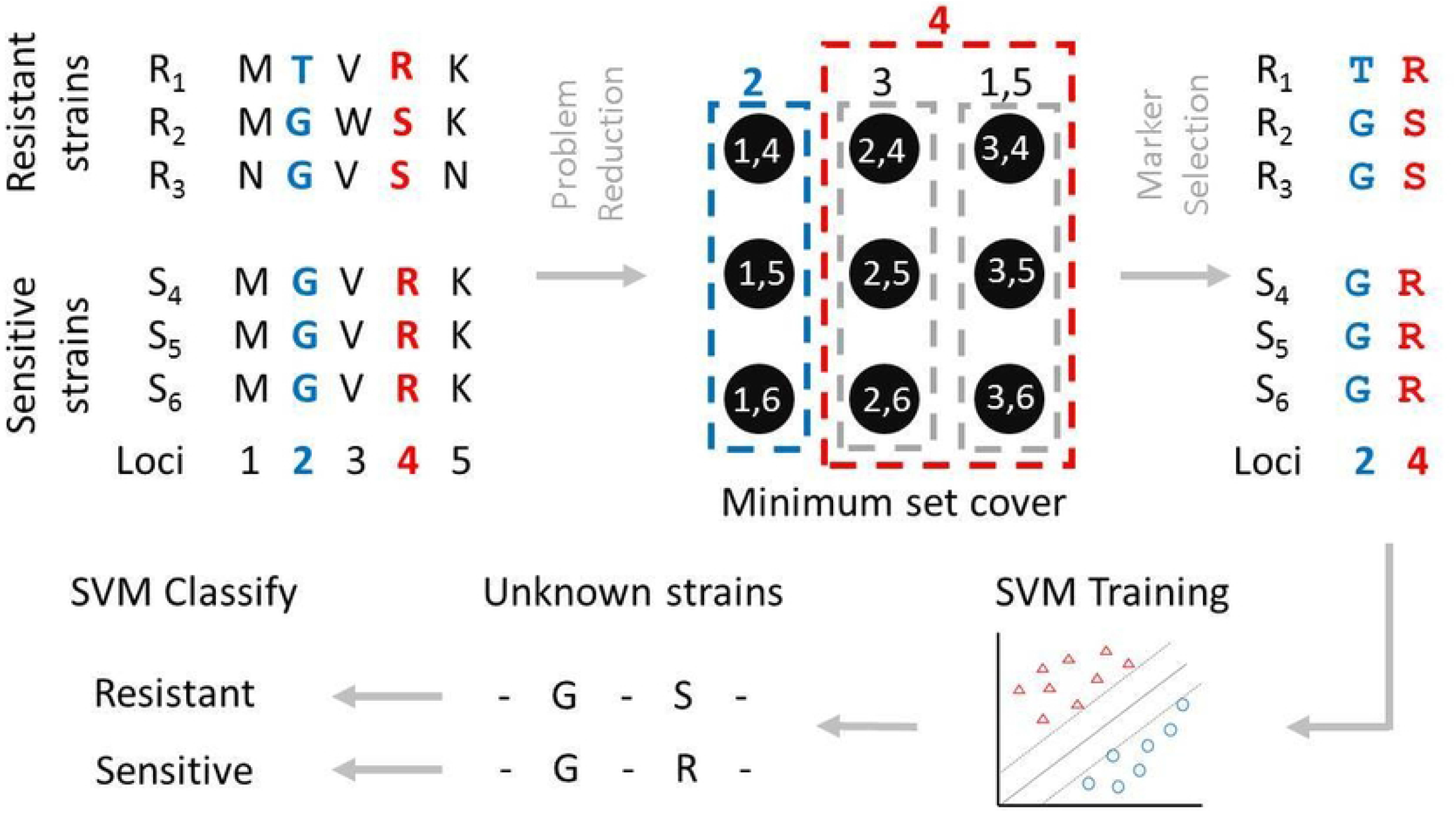
Illustration of combined mutations found by combinatorial optimization and machine learning. The markers are found by solving a minimum set cover problem. An SVM classifier is trained over the selected markers and used for prediction of unknown strains.

### Functional domains affected by combined mutations

The three combined mutations (PmrB451, PmrE168, PmrH292) located separately in PmrB, PmrE, and PmrH. PmrB belongs to the PmrCAB two-component system which produces PEtN attachment to lipid A. PmrE and PmrH are within the pathway of synthesizing L-Ara4N addition to LPS. To understand the functional impact of these mutations, the protein domains of PmrB, PmrE, and PmrH are investigated (Fig 3). Four and three protein domains are annotated within PmrB and PmrE, respectively, while PmrE is not annotated with any known domain. PmrB451 belongs to a histidine kinase-like ATPases domain (called HAPTase_c), which is responsible for activating PmrA via phosphorylation and for regulation of L-Ara4N synthesis and lipid A modification. On the other hand, PmrE168 is located within the UDP-GLC/GDP domain, which is the binding domain of UDP-glucose/GDP-mannose dehydrogenase essential for lipid A modification.

**Fig 3.**
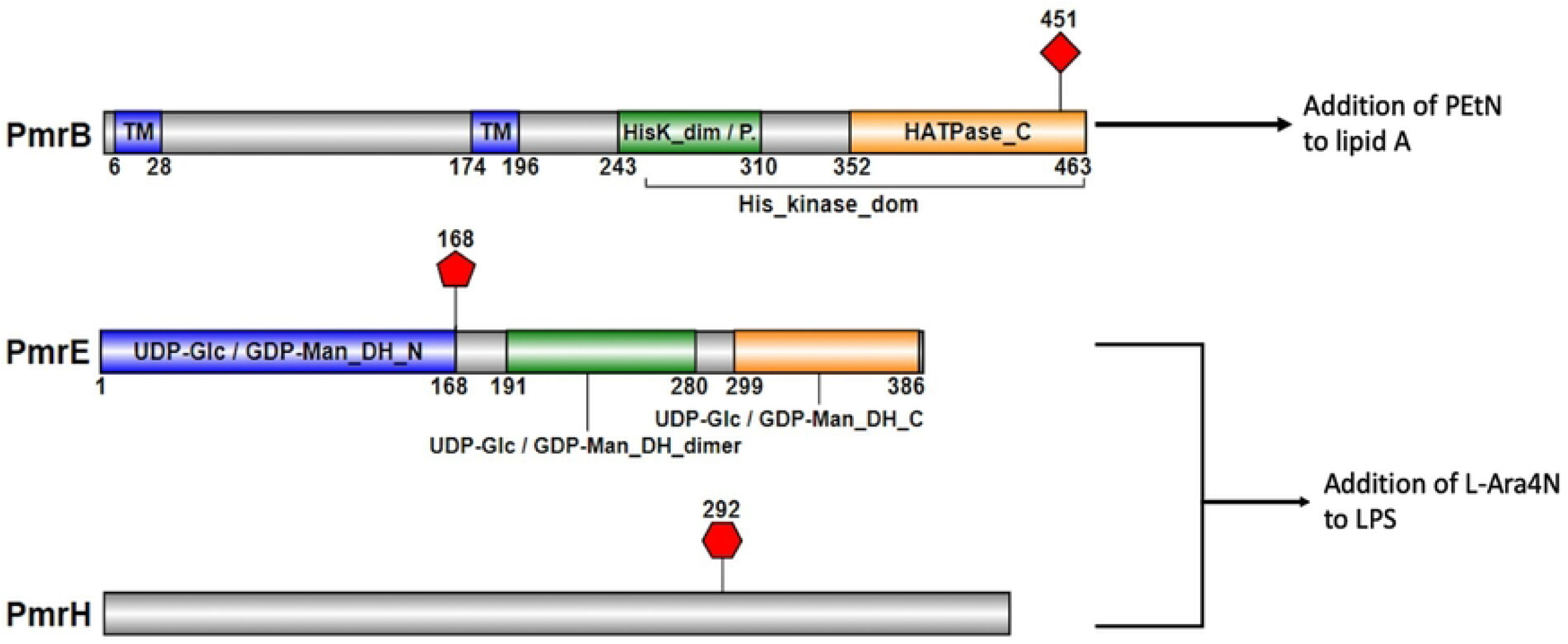
Functional domains impacted by PmrB451, PmrE168, and PmrH292 mutations. PmrB451 is within the histidine kinase-like ATPases domain. PmrE169 belongs to the UDP-GLC/GDP domains. PmrH292 is not annotated with any known domain.

## Discussion

The study compared the colistin-resistant mutations identified by single locus analysis and by combinatorial optimization. Each of the individual mutations identified by single locus analysis exhibited an association with statistical significance. However, their ability to accurately predict colistin resistance was still insufficient, even after combining partial or all identified mutations. As antibiotic resistance can be caused by multiple chromosomal mutations located on multiple genes, traditional statistical approaches were unable to discover combinations of multiple epistatic-interacted mutations. Furthermore, a combinatorial-optimization approach obtained a set of colistin-resistant mutations, and a combination of them showed a higher association. Similar observations were reported by other investigators. In *Mycobacterium tuberculosis*, mutations of *rpoB, katG, inhA* and *ahp* genes have shown increasing levels of multidrug resistance [24]. Mild effects of incidence mutations were usually more tolerable, while different structural effects brought by mutational combinations reflected differences in their functional importance [25].

Several studies have suggested that chromosomal resistance mutations often exhibit complex epistatic interactions among multiple genes [10, 26]. The fitness cost of resistance to a given antibiotic may also dependent on specific beneficial/deleterious alleles at other loci (i.e., genetic background) [12], and combination of them may sometimes even outcompete the sensitive strains [27]. To the best of our knowledge, most works studying epistatic interactions have only investigated a limited number of combined mutations [9], especially those involving gene knockout or editing experiments. This is because the number of possible combinations grows exponentially with respect to the number of mutational loci, and it is extremely difficult to know in advance which mutations interact epistatically. This work demonstrates a computational framework that can select possible epistatic-interacted mutations from hundreds of loci, which can facilitate future studies of epistatic interactions in a large set of genetic mutations.

Among the three combined mutations found, one mutation occurred in the histidine kinase-like ATPases domain in PmrB. PmrB is part of a two-component system that regulates the Pmr/Arn locus and PETn modification, which is a histidine kinase that activates PmrA by phosphorylation. Activated PmrA binds promoters within its regulone directly, increasing expression of L-Ara4N synthesis and modify lipid A [28]. Histidine kinase-like ATPases domain mutations have been reported to confer colistin resistance in *Salmonella enterica* serovar Typhimurium [29], *Acinetobacter baumannii* [30, 31] and *Pseudomonas aeruginosa* [32-34]. The fitness cost of colistin-resistant mutation on PmrB was conflicting in previous studies. During low dose colistin treatment from a patient with *Klebsiella pneumoniae* infection, a resistant mutation on PmrB was emerged without much fitness cost [35], as the resistant allele can be retained for up to 50 generations in the absence of colistin. On the other hand, a substantial fitness cost of colistin-resistant mutation on PmrB is required in *Acinetobacter baumannii* [4]. As the PmrB mutation coexists with other mutations (in PmrE and PmrH) in our study, this implies that a substantial amount of fitness cost is inevitable, at least in *S alage*.

The other PmrE mutation occurred in the UDP-glucose/GDP-mannose dehydrogenase (N-terminal). PmrE modify LPSs with 4-amino-4-deoxy-L-arabinose (L-Ara4N). In the presence of NAD^+^, PmrE oxidizes UDP-glucose (UDP-Glc) to UDP-glucuronic acid (UDP-GlcA) by UDP-Glc dehydrogenase, which is essential for lipid A modification with Ara4N and polymyxin resistance [36]. NAD binding domain mutation is associated with overexpression of PmrE leading to colistin resistance [5]. Interestingly, PmrE and PmrB mutations have been known to highly correlate with colistin resistance in *Pseudomonas aeruginosa* [5], where mutations in PmrB tend to emerge first, followed by mutations in PmrE. Our computational approach identified these two epistatic-interacted mutations from hundreds of loci.

A novel mutation in PmrH was discovered by this computational approach. PmrH (or arnB) encoding uridine 5′-(beta-1-threo-pentapyranosyl-4-ulose diphosphate) aminotransferase plays an important role in colistin resistance [37]. Deletion of entire PmrH was shown to develop colistin resistance through the addition of l-Ara4N to lipid A. However, no literature on amino acid substitution level was conducted on this gene, although our results suggest it is involved in increasing resistance to colistin. This study confirmed previous *Pseudomonas aeruginosa* study that resistance to colistin of combined mutations (from 16 genetic background) is indeed greater than that of individual mutations [13]. As the number of mutational combinations grows exponentially with respect to the number of loci, most previous works can only validate a subset of them. Our proposed computational approach systematically identifies epistatic-interacted mutations from a large number of potential loci within a reasonable period of time.

In the study we compare single locus analysis using single locus analysis to the identification of multiple combined mutations using novel combinatorial approaches. Single locus analysis identified significant associations, but with low prediction accuracies. However, the combination of combinatorial and machine learning approaches identified three mutations and achieved significantly higher prediction accuracies.

The genetic and molecular mechanisms underlying colistin resistance in *S. algae* are poorly understood. In many cases, chromosomal mutations interact epistatically to lower the overall fitness cost of resistance to antibiotics. These complex epistatic interactions among multiple genes bring great challenges. We integrated combinatorial and machine learning to compare the analysis of a single locus with the analysis of multiple combined mutations and identified a combination of three mutations that could achieve a much higher level of accuracy. We demonstrated an effective computational approach for identifying epistatic-interacted mutations from a large number of loci in this study.

## Acknowledgments

None.

## Data availability statements

The sequencing data is available in the NCBI GenBank under BioProject accession number PRJNA312015.

## Supporting information

**S1 Fig. The ROC curves**. Individual mutations are compared with combined mutations found by different approaches.

